# Non-parametric Functional Muscle Network as a Robust Biomarker of Fatigue

**DOI:** 10.1101/2021.09.29.462080

**Authors:** Rory O’Keeffe, Seyed Yahya Shirazi, Jinghui Yang, Sarmad Mehrdad, Smita Rao, S. Farokh Atashzar

## Abstract

The possibility of muscle fatigue detection using surface electromyography has been explored and multiple biomarkers, such as median frequency, have been suggested. However, there are contradictory reports in the literature which results in an inconsistent understanding of the biomarkers of fatigue. Thus, there is an unmet need for a statistically robust sEMG-based biomarker for fatigue detection. This paper, for the first time, demonstrates the superior capability of a non-parametric muscle network to reliably detect fatigue-related changes. Seven healthy volunteers completed a lower limb exercise protocol, which consisted of 30s of a sit-to-stand exercise before and after the completion of fatiguing leg press sets. A non-parametric muscle network was constructed, using Spearman’s power correlation and showed a very reliable decrease in network metrics associated with fatigue (degree, weighted clustering coefficient (WCC)). The network metrics displayed a significant decrease at the group level (*degree, WCC*: *p <* 0.001), individual subject level (*degree*: *p <* 0.035 *WCC*: *p <* 0.004) and particular muscle level (*degree*: *p <* 0.017). Regarding the decrease in mean degree connectivity at particular muscles, all seven subjects followed the group trend. In contrast to the robust results achieved by the proposed non-parametric muscle network, classical spectrotemporal measurements showed heterogeneous trends at the particular muscle and individual subject levels. Thus, this paper for the first time shows that non-parametric muscle network is a reliable biomarker of fatigue and could be used in a broad range of applications.

## I. Introduction

**M**USCLE fatigue is the decrease of contractile force generation ability and challenges the motor control and performance for task completion [1]. Fatigue is a physiological phenomenon which has both positive and negative effects. On the positive side, fatigue has been postulated to drive muscle adaptation [2] and exercise-induced hypoalgesia [3]. On the negative side, fatigue is a prevalent source of injury in sports and non-sport activity both to the muscles and to the other structures of the skeletal system [4], [5], [6], [7], which negatively affects the person’s performance in their working environment and daily lives. Individuals with motor impairments (such as those associated with stroke) and surgical interventions (such as those associated with knee ligament repair) may experience an altered pattern of muscle fatigue, likely because muscle recruitment also changes with these impairments and interventions [8], [9], [10]. Therefore, the objective quantification of fatigue is essential for both injury prevention and monitoring the rehabilitation and intervention process.

The human motor control in the central and peripheral nervous systems responds to muscle fatigue by altering the normal movement biomechanics [11], [12], changing the muscle groups required for low-intensity tasks [13], increasing cocontraction [14] and likely modulating the contralateral limb muscle activation either as a compensatory mechanism or due to the neurophysiological and biochemical interlinkage [15], [16], [17].

Muscle fatigue can be categorized as either maximal or sub-maximal, depending on the activity which has caused the fatigue. In sub-maximal activity, the subject can increase motor unit recruitment to counteract fatigue’s effect of force reduction [18]. On the contrary, all available motor units are being recruited during maximal activity, so a subject must alter firing patterns to counteract the force reduction effect. Due to these differences, sub-maximal fatigue has a slower onset time at both the peripheral and central levels. Sub-maximal muscle fatigue can be induced by moderate intensity resistance training, and is highly clinically relevant to patients’ symptoms during rehabilitation [19].

The electromyographic changes in the exercised muscle, which stem from changes in muscle recruitment process, are usually observable as a decrease in median frequency (MDF) [20]. Additionally, changes in root mean square (RMS) of the sEMG in the exercised muscle have been reported, although the nature of these changes is inconsistent across the relevant studies, for example [21], [22], as variability between anatomy types and muscle groups can affect the results. However, fatigue-related changes are not confined to the exercised muscle alone [23].

Another common sEMG analysis method involves computing muscle synergies, i.e., low-dimensional activations that the nervous system can combine to produce a complex movement [24]. Muscle synergies show retained synergistic activation before and after fatigue but with different amplitudes, which suggests that the main control is on the activation amount of each synergy, and the muscle groups themselves may not change because of fatigue [25]. However, based on the literature, it is known that the patterns of changes regarding synergy measures are specific to each person, and their muscle recruitment history and may not respond in a consistent manner to fatigue [26], [27], [28], [29], [30]. Therefore, muscle synergy analysis is susceptible to inter-subject variability which has already resulted in a heterogeneous literature regarding its response as a biomarker of fatigue. Thus it might not be a robust and reproducible method to monitor fatigue. The non-consistent changes in muscle activity patterns measured using the aforementioned metrics necessitates designing a more complex method that can potentially provide the homogeneous and consistent group-level and individual-level results which are needed as a trustworthy “biomarker”.

A new candidate can be functional muscle connectivity network which has attracted researchers during the last few years due to the corresponding holistic network view on the synchronicity of muscle recruitment during functional tasks. It should be noted that a functional muscle connectivity network is a new way of understanding how the central nervous system (CNS) is controlling synchronicity between muscles and distributing the activation to conduct functional tasks [31], [32]. Thus, the muscle network is considered to reflect both peripheral and central nervous changes, which is also imperative for fatigue.

The concept of a functional muscle connectivity network has been recently investigated for non-fatiguing normal tasks and shown high sensitivity to subtle motor changes [33], [34], [31]. This level of task sensitivity to changes in (non-fatiguing) motor tasks has not been reported before using muscle synergy analysis or other forms of conventional spectrotemporal metrics. This also motivates the investigation of the utilization of functional muscle network for the assessment of fatigue. However, it should be noted that all existing literature on functional muscle network has focused on intermuscular coherence analysis (to construct the network), which is a linear analysis in the spectral domain. However, intermuscular coherence has also shown inconsistent trends during fatigue. For example, while some muscle pairs show decreased coherence or interactions from before to after the fatigue, other muscle pairs may not change or even increase their coherence after the fatigue process [35], [36]. The aforementioned issue can be explained by the fact that neural synchrony changes in a non-linear manner [37].

Thus, a non-parametric analysis of coupling may provide a more transparent understanding of the neurophysiological changes caused by fatigue. In response to the above-mentioned heterogeneous literature, in this work we will investigate the efficacy of a non-parametric and non-linear functional muscle connectivity network which can highlight monotonic but non-linear synchronicity in the activations of muscles. This paper hypothesizes that physiological fatigue can be readily detected by a non-parametric form of intermuscular connectivity. Unlike conventional connectivity metrics which are linear, such as Pearson’s correlation or coherence, a non-parametric technique can detect complex changes in central and peripheral nervous systems and is hence utilized in this work.

The purpose of this study is to quantify non-parametric functional muscle network changes in a sit-to-stand task due to unilateral leg-press exercise. The leg press exercise would mostly fatigue rectus femoris and vastus lateralis muscles, since quadricep muscles are known to be most active in knee extension [38]. Therefore, we hypothesize to observe the most changes in the non-parametric muscle network in these two muscles during the sit-to-stand task. The choice of sit-to-stand task is supported by the literature which uses this task as a simple test of composite lower extremity muscle strength [39]. The results of this paper show that not only the group averages are strong indicators of fatigue, especially for vastus lateralis and rectus femoris, but also the individual levels indicate similar trends for all the participants, suggesting that functional muscle networks can be indeed effective and robust biomarkers for fatigue.

## II. Methods

Seven healthy subjects (three females, four males) with a mean (SD) age of 30.4 (6.4) years and a mean (SD) BMI of 22.95 (3.6) *kg*.*m*^2^ participated in the study. The institutional review board of New York University approved the study and subjects provided their written consent after they received the study description. Subjects did not report any lower limb injury or impairment at the time of recording.

### A. Experimental Procedure

Subjects performed a sit-to-stand task before and after submaximal fatigue was induced by a single-session of moderate intensity resistance training (Fig. 1a). The 30 second sit-to-stand task involved repeatedly standing up from a seated position in a chair, for 30 seconds [40], [41]. After performing the first set of sit-so-stand, sub-maximal fatigue was induced in the subjects by performing four sets of resistance exercises with a leg press machine, unilaterally [42], [43]. The subject’s one repetition maximum (1RM) was determined first. Subjects then performed four sets at 50% of the 1RM, with a target of 30 repetitions in the first set and a target of 15 repetitions for the last three sets. The last set was specifically performed to failure, defined as the inability to complete a repetition. The subject-wise number of repetitions for each set are shown in Table I. Subjects were given a fixed recovery time (30 seconds) between sets. Ninety seconds after completing the last repetition of the leg press, subjects performed the second set of sit-to-stand. Subjects were instructed to complete the sit-to-stand at a natural rate, and to maintain that rate for both sets. As illustrated in Table II, the number of repetitions was comparable before and after fatigue, for all subjects.

**Fig. 1.**
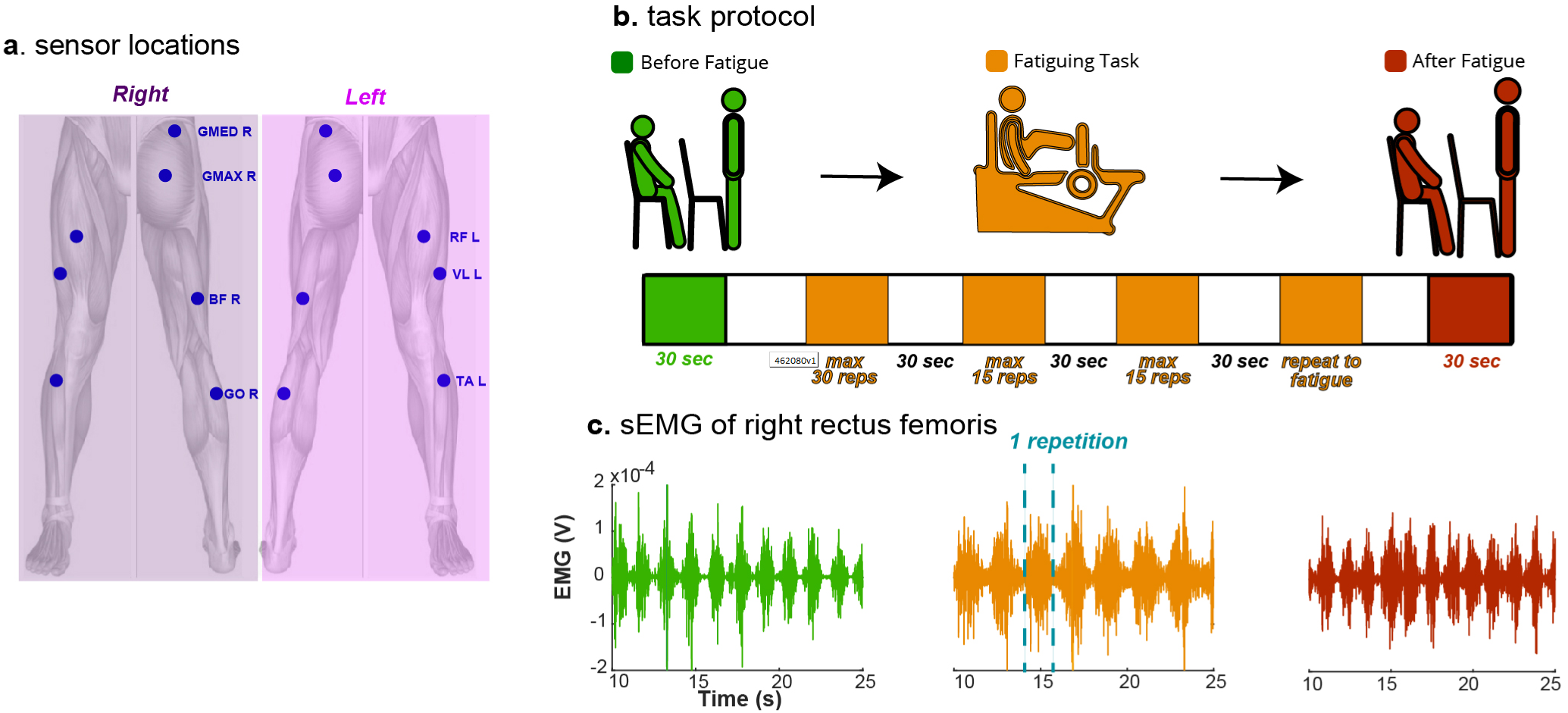
Experimental Outline. (a) Seven bipolar sEMG sensors were placed bilaterally on the anterior and posterior leg muscles. GMED = Gluteus Medius, GMAX = Gluteus Maximus, BF = Biceps Femoris, GO = Gastrocnemius, RF = Rectus Femoris, VL = Vastus Lateralis, TA = Tibialis Anterior. (b) Subjects performed 30s of sit-to-stand repetitions before and after a fatiguing task. The fatiguing task consisted of four sets of resistance exercises with a leg press machine. (c) A sample subject’s sEMG recording from RF during the sit-to-stand pre-fatigue, fatiguing exercise and sit-to-stand post-fatigue.

**TABLE I.** Leg Press Repetitions By Set

| Subject | Set #1 | Set #2 | Set #3 | Set #4 |
| --- | --- | --- | --- | --- |
| 0 | 30 | 15 | 12 | 12 |
| 1 | 30 | 15 | 12 | 11 |
| 2 | 30 | 15 | 15 | 15 |
| 3 | 30 | 15 | 15 | 15 |
| 4 | 30 | 15 | 15 | 15 |
| 5 | 27 | 12 | 9 | 7 |
| 6 | 30 | 15 | 10 | 7 |
| Mean | 29.57 | 14.57 | 12.57 | 11.71 |
| Std | 1.13 | 1.13 | 2.51 | 3.59 |

**TABLE II.** Sit-to-stand Repetitions, Before and After Fatigue

| Subject | Before | After |
| --- | --- | --- |
| 1 | 22 | 20 |
| 2 | 21 | 24 |
| 3 | 12 | 12 |
| 4 | 10 | 11 |
| 5 | 17 | 19 |
| 6 | 15 | 17 |
| 7 | 10 | 11 |

sEMG signals were recorded from fourteen sensors (Fig. 1b), using the wireless Trigno sEMG system (Delsys Inc., Natick, MA), with a sampling frequency of 1259 Hz. Fourteen bipolar Trigno Avanti sensors were used bilaterally for (i) Rectus Femoris (RF), (ii) Vastus Lateralis (VL), (iii) Tibialis Anterior (TA), (iv) Gluteus Medius (GMED), (v) Gluteus Maximus (GMAX), (vi) Biceps Femoris (BF) and (vii) Gastrocnemius (GO). The skin surface was thoroughly wiped prior to sensor placement. Sensors were placed parallel to the direction of the muscles. Following the recording, signals were pre-processed using MATLAB R2020b (MathWorks Inc., Natick, MA). The first and last one second of all trials were clipped out. Then, a zero phase Butterworth high pass filter at 1 Hz was applied.

### B. Non-parametric Muscle Network

A zero-phase Butterworth low pass filter was applied at 50 Hz, such that the resultant sEMG signals were in the 1-50 Hz range. Spearman power correlation networks were constructed for before and after fatigue. Spearman power correlation (*ρ*_*xy*_) between two sEMG signals *x*(*t*) and *y*(*t*) was computed. The power time series *x*^2^(*t*), *y*^2^(*t*) were first calculated and then each power time series was rank transformed. For example, an sEMG signal with *n* samples will have its power values replaced by ranks from 1 to *n*, in ascending order - the maximum power value will be assigned the rank *n*. After rank transforming *x*^2^(*t*) to *p*_*x*_(*t*) and *y*^2^(*t*) to *p*_*y*_(*t*), *ρ*_*xy*_ is computed according to:

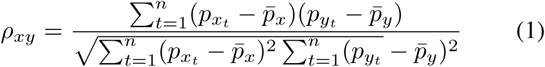

For the muscle network, the magnitude |*ρ*_*xy*_| was computed between each sensor pair, across the sit-to-stand trial duration. Using the magnitude |*ρ*_*xy*_| interprets a negative Spearman power correlation between *x*(*t*) and *y*(*t*) of − *ρ*_*xy*_ as having equivalent connectivity to the positive correlation *ρ*_*xy*_. Each node in the network represents a muscle, and the width of each line illustrates |*ρ*_*xy*_|. The degree of each node, *D*_*i*_, is the average of all edges connected to the node. If the muscle network is represented by adjacency matrix *A, D*_*i*_ is defined as:

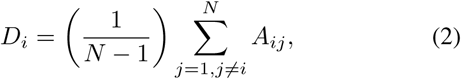

where *N* is the number of nodes. A node’s weighted clustering coefficient (*WCC*_*i*_) gives the measure of how well that node is connected to its neighbors. The weighted clustering coefficient is defined as:

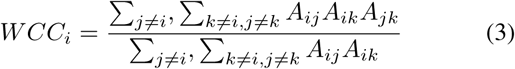

A node which is not connected to its neighbors will have *WCC*_*i*_ = 0, while a node which is very well connected to its neighbors has *WCC*_*i*_ = 1.

### C. Conventional Spectrotemporal Measurements

To compare the proposed network results to the existing attempted biomarker methods, RMS, power spectral density (PSD) and median frequency (MDF) were computed for particular muscles. All measurements were calculated by considering the full trial duration, before and after fatigue, for each subject. After low-pass filtering the signal at 50 Hz (resultant band: 1-50 Hz), RMS and PSD were calculated. In the case of PSD, the median PSD in the 1-50 Hz range was used. Finally, the median frequency was computed on a signal which had been low-pass filtered at 400 Hz (resultant band: 1-400 Hz), with zero-phase notch filters (half-width = 2.5 Hz) at multiples of 60 Hz.

### D. Statistical Analysis

To evaluate the statistical trends observed in absolute power correlation muscle networks, (i) group-level, (ii) individual subject and (iii) particular node analyzes were conducted. In all cases, connectivity (*ρ*_*xy*_) was evaulated at each of the fourteen nodes using degree and WCC. A group connectivity distribution (Fig. 3) which includes all subjects contains *n* = 14 ×7 = 98 samples, while an individual subject distribution has *n* = 14 (Fig. 4). When considering the connectivity statistics of particular nodes (Fig. 5), the node’s absolute power correlations for all subjects are included (*n* = 7). Similarly, when considering the spectrotemporal measurements (e.g. RMS), a given muscle’s value for all subjects (*n* = 7) is used for the statistical analysis.

The Kolmogorov-Smirnov test for normality rejected the normal distribution hypothesis for the absolute Spearman’s power correlation distributions. The Wilcoxon signed-rank test was used to test statistical significance, with the significance level *α* = 0.05. For the sake of even comparison, the Wilcoxon signed-rank test was also utilized for measuring significance of spectrotemporal measurement distributions.

## III. Results

### A. Group Analysis of Non-parametric Muscle Network

To test the subjects’ response to fatigue, group level analysis of the non-parametric muscle network was conducted. The group-level results show a strong trend of decreasing lower limb network connectivity from before to after fatigue. The subject-mean network (i.e., the mean network calculated by the average of each pairwise connection across the subjects) is illustrated in a heat map (Fig. 2a) and a network map (Fig. 2b) forms. For the mean networks, most of the connectivity edges show a fatigue-induced decrease. To confirm these trends, graph theory metrics of connectivity (degree, WCC and global efficiency) were computed and evaluated in a statistical analysis. Group-level statistical analysis of all subjects’ network metrics confirms that non-parametric intermuscular connectivity significantly decreases from before to after fatigue (Fig. 3). All network metrics were significantly higher before the fatiguing exercise than after (*degree and WCC: p* < 0.001, *global efficiency: p* = 0.016). These results illustrate that the non-parametric muscle network can detect the effect of fatigue at the group level.

**Fig. 2.**
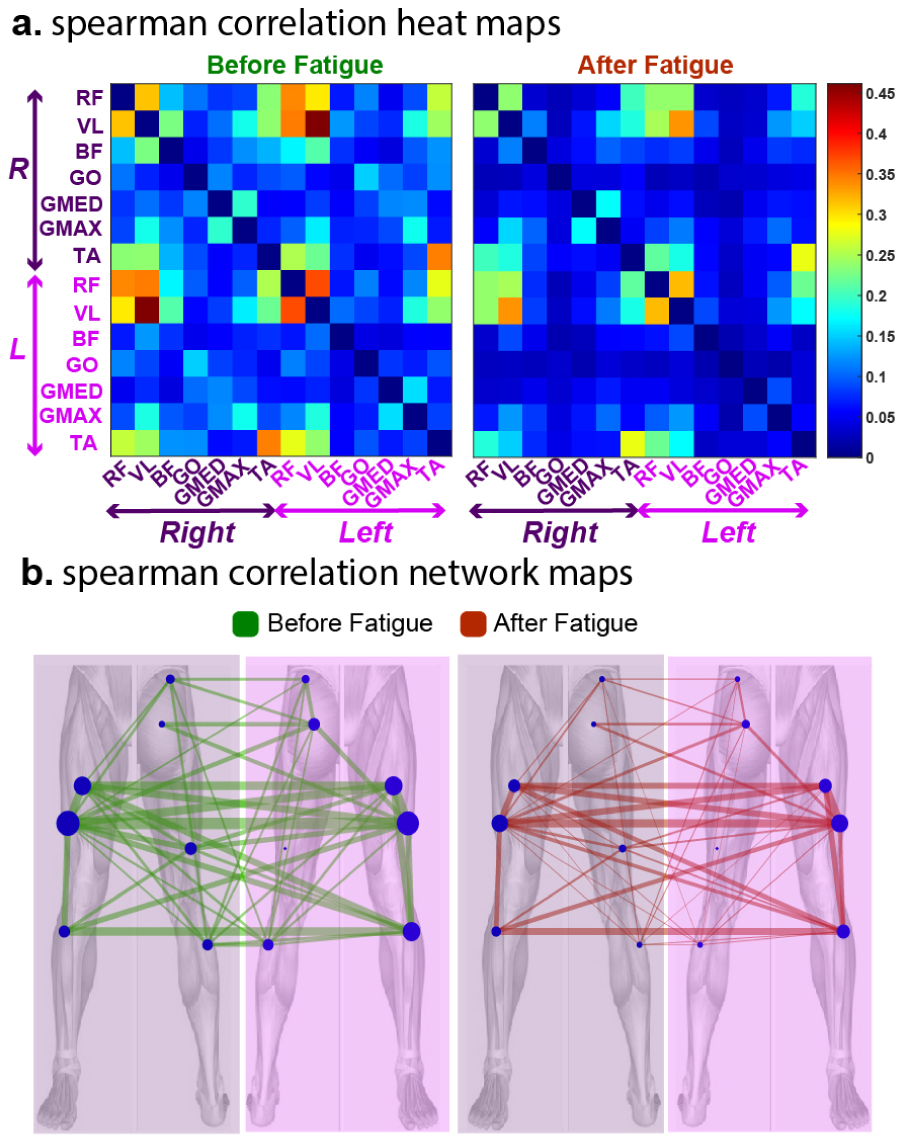
Subject-mean absolute Spearman’s power correlation network. The mean network across all subjects was computed for before and after fatigue. **a**. The heat map illustrates the changes from before to after, for all network edges. **b**. The top 50% most changeable edges from before to after fatigue are shown in the network map.

**Fig. 3.**
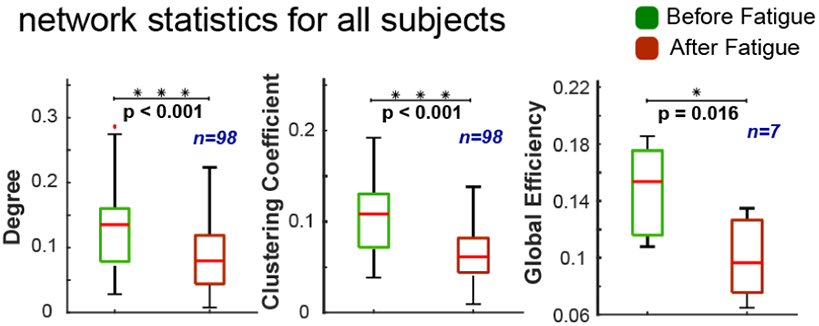
Statistical analysis for all subjects, comparing the non-parametric muscle network changes from before to after fatigue. All fourteen nodes for all subjects are included in each distribution for degree and WCC (*n* = 98). Each single global efficiency value for all subjects’ networks are included in the global efficiency distributions (*n* = 7). The effect of fatigue was investigated by testing for statistical significance between the distributions by using the Wilcoxon signed-rank test. For all network metrics, the null hypothesis that the distributions are similar was rejected at the 0.05 significance level *(degree and WCC: p* < 0.001, *global efficiency: p* = 0.016).

The effect of decreasing connectivity with fatigue appears to be most pronounced in the quadricep muscles (Fig 2). In Fig 2b, we note the visible decrease in the size of the circles (proportional to degree of the node) for the two quadricep muscles, i.e. RF and VL, in a bilateral manner. In Fig. 2a, it can be observed that a high connectivity between ipsilateral RF and VL exists on both left and right sides. In both cases, there appears to be a marked decrease from before to after fatigue. From Fig. 2b, it can be also observed that a high connectivity exists for contralateral muscle pairs, for example left RF with right RF, left VL with right VL. These two connections also appear to show a decrease from before to after fatigue. The contralateral effect (will be discussed later) highlights the systematic effect of fatigue on neural aspects of motor control in tasks which would require bilateral synchronicity/coordination, or it can be caused by a compensatory mechanism.

### B. Individual Subject Analysis of Non-parametric Muscle Network

A robust biomarker of fatigue will be responsive at both the group and individual levels, hence non-parametric muscle network analysis was undertaken for each subject. All subjects showed the pattern of decreasing connectivity from before to after fatigue. This is illustrated in Fig. 4 - where the statistics of the full network metrics (node degree, node WCC, network global efficiency) for each subject are shown. At the individual level, the effect of fatigue is illustrated by a statistically significant decrease from before to after for both network metric distributions (*degree: p ≤* 0.035, *WCC: p ≤* 0.004) and a consistently decreasing network global efficiency. This subject-specific result further emphasizes the robustness of the non-parametric muscle network’s response to fatigue.

**Fig. 4.**
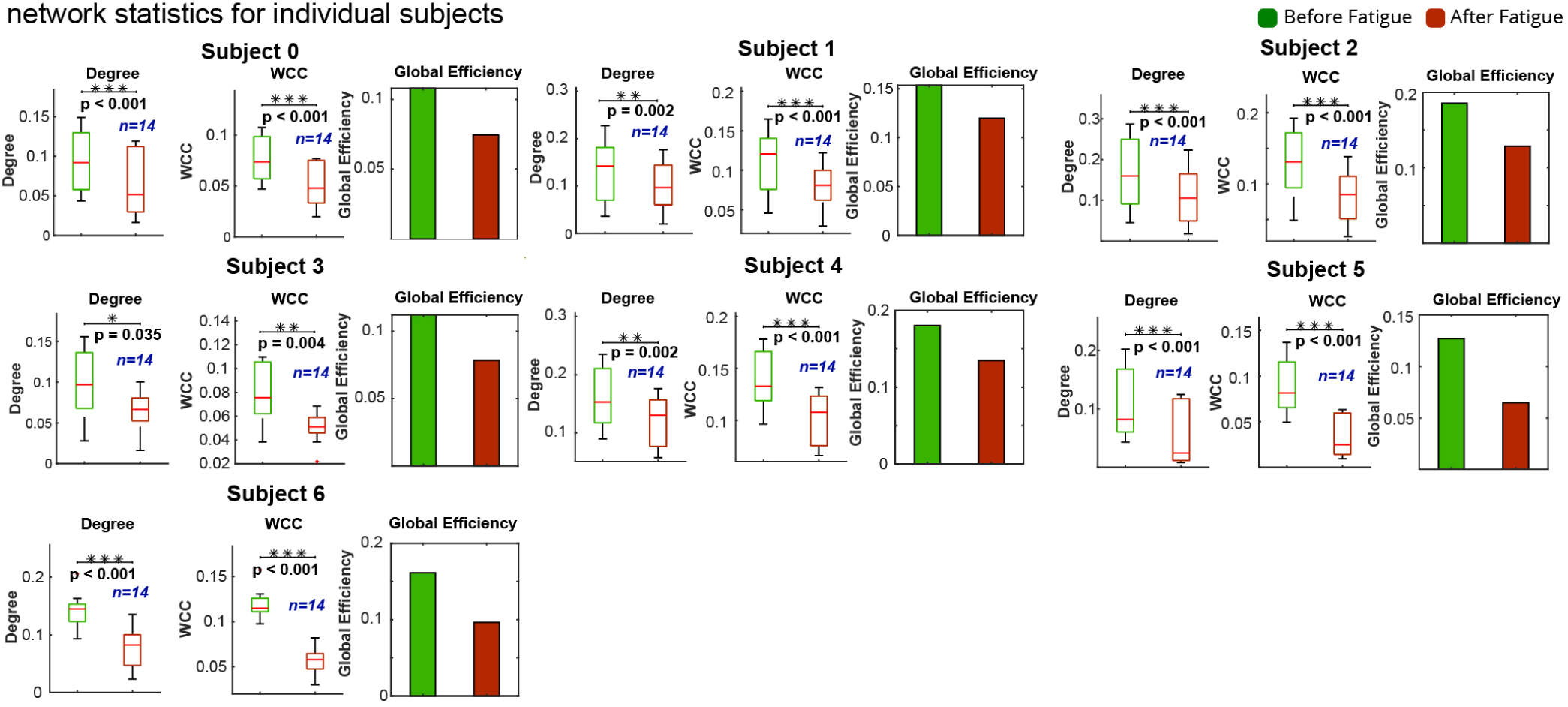
Non-parametric muscle network statistics for each subject, comparing before and after fatigue. The degree and WCC for all fourteen nodes form the distributions. The effect of fatigue was investigated by testing for statistical significance between the distributions by using the Wilcoxon signed-rank test. For all subjects, the null hypothesis that the distributions are similar was rejected at the 0.05 significance level *(degree: p* < 0.035, *WCC: p* < 0.004).

### C. Nodewise Analysis of Non-parametric Muscle Network

To identify the location with the most consistent response to fatigue, the non-parametric connectivity of particular nodes (muscles) was analyzed. Across all the subjects, the selected muscles (RF, VL, GO and TA) showed a consistent decrease in degree after fatigue. This is illustrated in Fig. 5, where for each muscle, the distribution across all subjects is shown, for both left and right legs. For each muscle, there is a significant decrease in degree (*p=0*.*016*). This demonstrates that not only does the overall network show a significant decrease from before to after fatigue, individual nodes also exhibit the connectivity decline. Moreover, the blue hairlines show the individual subject changes, and this most strongly emphasizes the robustness of the result, since all subjects follow the group trend of decreasing connectivity.

**Fig. 5.**
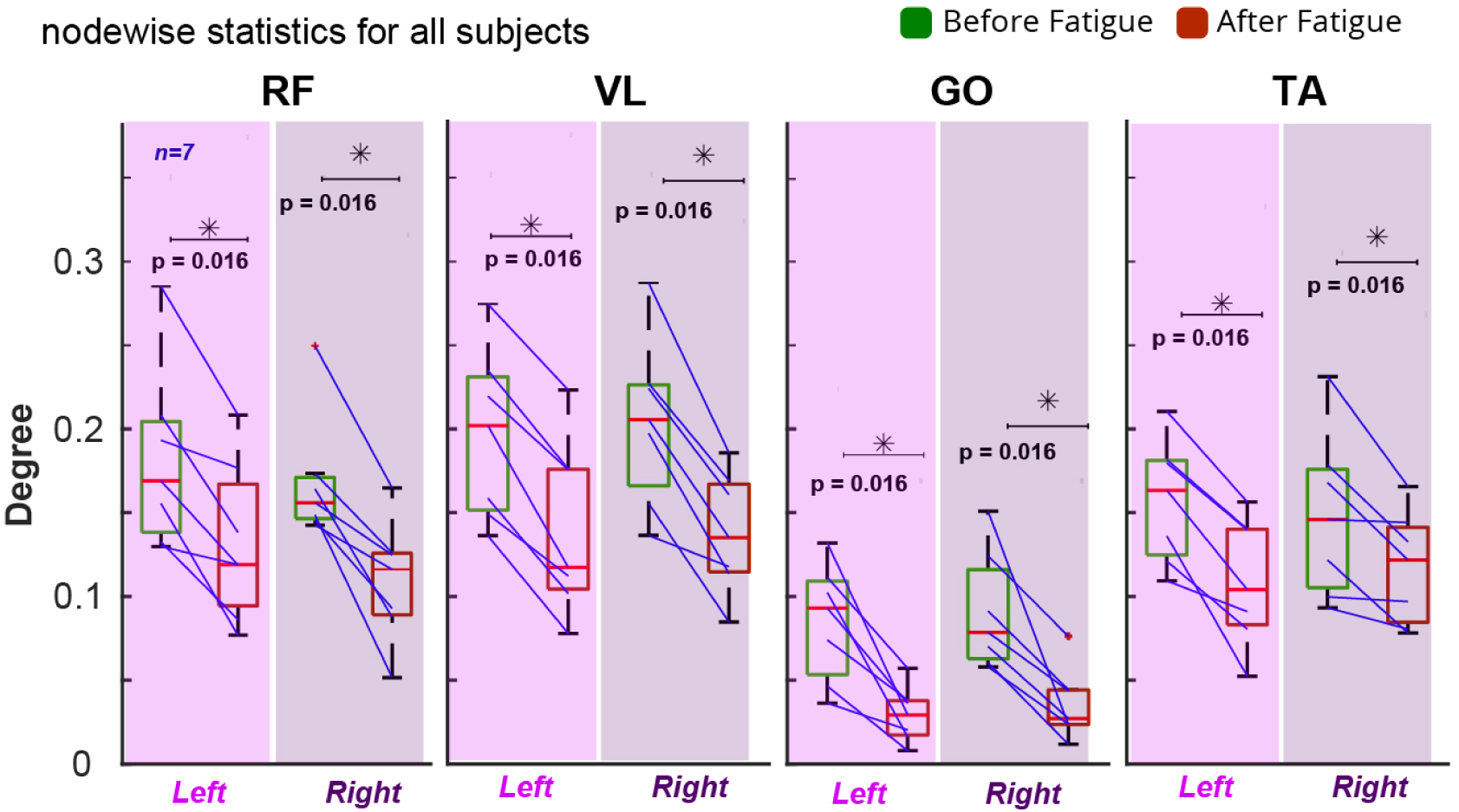
Particular muscle (nodewise) non-parametric connectivity statistics across all subjects, comparing before and after fatigue (*n* = 7). The blue lines show the change in degree from before to after fatigue for each subject. The selected muscles are rectus femoris (RF), vastus lateralis (VL), gastrocnemius (GO) and tibialis anterior (TA). For all four selected muscles, the degree of all subjects decreased from pre to post-fatigue. The consistent decrease in network degree was observed on both the left and right sides (both sides: *Wilcoxon signed rank test p* = 0.016).

### D. Comparison with Spectrotemporal Measurements

In contrast with the robust and consistent results obtained for the non-parametric intermuscular coupling in response to fatigue, classical sEMG-based spectrotemporal measurements of fatigue showed heterogeneous responses. The RMS, PSD (median in 1-50 Hz range) and median frequency for each muscle are shown in Fig. 6. In Fig. 6(a), there is no significant decrease in any muscle’s RMS from before to after resistance-training induced muscle fatigue, on either the left or right leg. In Fig. 6(b), there is an increase of PSD for right VL, while in Fig. 6(c), there is an increase in MDF for left GO. Overall, no consistent pattern of an increasing or decreasing spectrotemporal measurement was observed for the muscles shown, in sharp contrast to the non-parametric muscle network results.

**Fig. 6.**
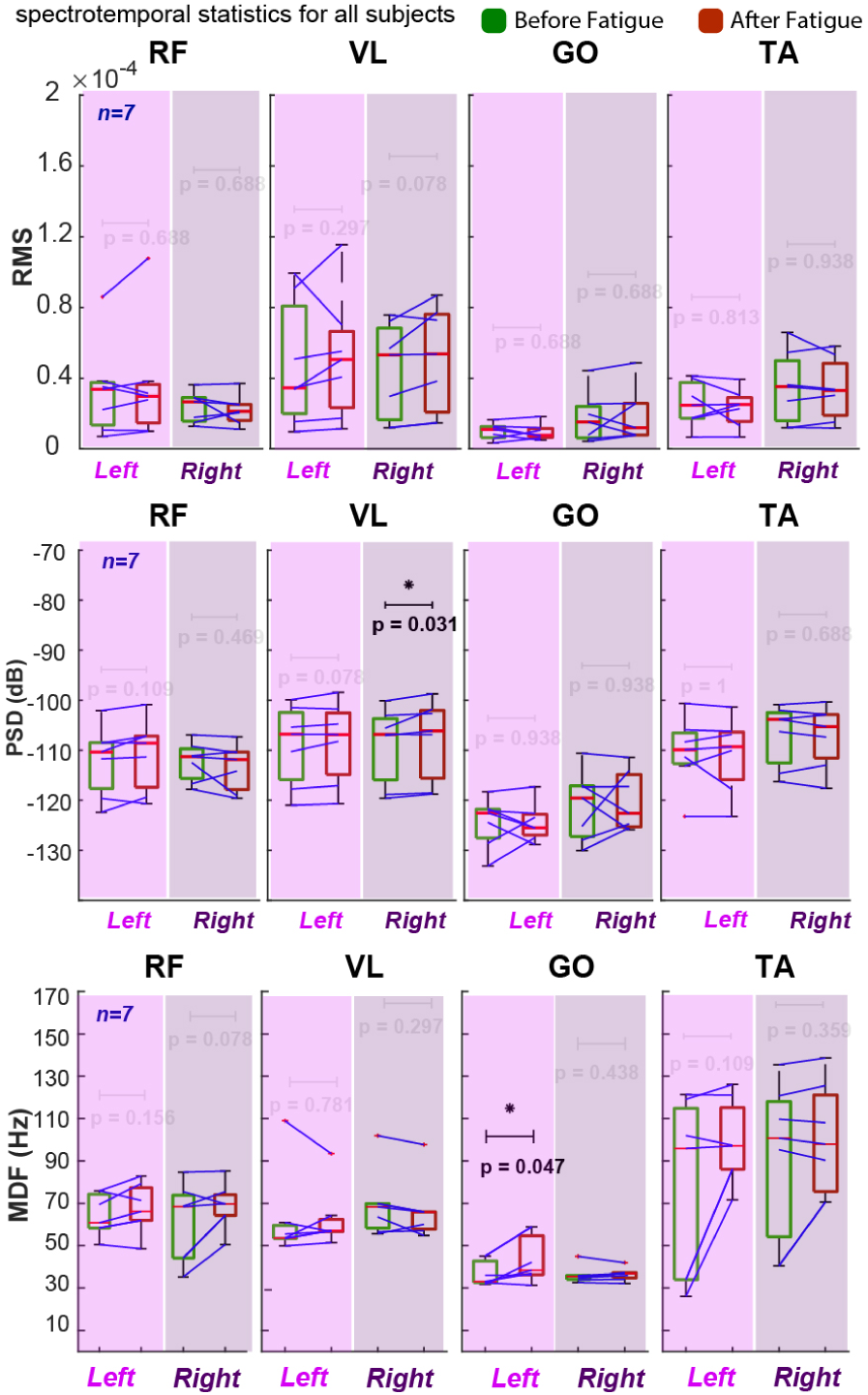
Statistical analysis of spectrotemporal measurements for particular muscles. Distributions are constructed for each muscle using all subject values for that measurement (*n* = 7). Blue hairlines show the individual subject changes. The selected muscles are rectus femoris (RF), vastus lateralis (VL), gastrocnemius (GO) and tibialis anterior (TA). (a) The change in RMS on both left and right is analyzed, before and after fatigue. No significant change from before to after fatigue is observed for left or right. Hairlines show inconsistent patterns. (b) Change in PSD (median in 1-50 Hz range), before and after fatigue. Right VL shows a small increase (*p* = 0.031). No other significant changes in PSD were observed, and hairlines show inconsistent patterns. (c) Change in MDF, before and after fatigue. Left GO shows a small increase (*p* = 0.047). No other significant changes in MDF were observed, and hairlines show inconsistent patterns.

## IV. Discussion

The non-parametric muscle network was demonstrated to be a statistically robust biomarker of fatigue, since it efficiently responded to fatigue-related changes at the group and individual levels. Moreover, key muscles were shown to have decrease in their degree for all subjects, emphasizing the ability of the network to detect degradation of the peripheral nervous system (PNS) response to central nervous system (CNS) command after resistance-training induced muscle fatigue. Additionally, conventional methods (RMS, PSD and MDF) showed heterogeneous responses to fatigue, highlighting the relative strength of the proposed method. The reliability of the proposed method can be utilized in settings such as physiotherapy and rehabilitation, where accurate quantification of fatigue is needed.

As hypothesized, the non-parametric muscle network was able to detect muscle fatigue. This is demonstrated by significant post-fatigue decreases in network metrics at the group (Fig. 3) and individual (Fig. 4) levels. The absolute Spearman’s power correlation detects fatigue-related decreases in the non-parametric coupling between muscle activations (in power form), which is illustrated by the network degree declining for all subjects (Fig. 4). The connectivity between adjacent nodes is reduced as shown by the WCC decreasing for all subjects (Fig. 4). The overall network communication is also diminished, with global efficiency consistently decreasing (Fig. 4). The fatigue-related degradation in the non-parametric intermuscular coupling was highlighted in four selected muscles (Fig. 5), and indeed all subjects showed a decrease for each muscle.

A secondary hypothesis was that the non-parametric functional muscle network could detect physiological muscle fatigue most particularly in the quadricep muscles, since they are most active during leg press and play a key role in the sit-to-stand. Examining the mean non-parametric muscle network in Fig. 2 showed a visible decrease in ipsilateral RF-VL bilaterally, and in contralateral RF-RF and VL-VL. Moreover, the hypothesis was thoroughly validated in Fig. 5, where all subjects showed a decrease in mean connectivity for RF and VL.

Additionally, two other key muscles which illustrated fatigue-related network change were observed to be GO and TA. This observation is in spite of the fact that these distal muscles of lower limb were not directly targeted by the fatiguing task. The change in TA can be attributed to a compensatory mechanism; since RF and VL were fatigued, TA (which is active during sit-to-stand, Fig. 6(b)) may need to provide supplementary, control, which could result in deviations from the typical network. In addition, the GO muscle showed a consistent decrease in degree after fatigue, for all subjects (Fig. 5). An important component of the uniform change in average GO connectivity is its non-parametric coupling with GMAX and GMED. Indeed, ipsilateral GO-GMED and ipsilateral GO-GMAX decreased post-fatigue for 7/7 and 6/7 subjects respectively on the fatigued side. These results indicate that non-parametric synergistic proximal-distal coupling is diminished after resistance-training induced fatigue. It should be noted that fatigue has been previously shown to effect lower limb joint coupling [44]; thus the reduced hip-ankle muscular coupling here is perhaps indicative of a fatigue-related decrease in the precision of lower limb motor control.

The results support the notion that the disruption of motor control, induced by fatigue, can result in a systematic response of the proposed non-parametric muscle network. The muscle network response is influenced by subtle functional motor changes and the overall synchronicity of PNS with commands from CNS. The decrease in the connectivity at the fatigued side (shown for the first time in this paper) (Fig. 5), can be explained as the reduction in responsiveness of the fatigued muscles to CNS commands. The decrease of the connectivity at the contralateral side (reported in Fig. 5) can be due to compensatory mechanisms, which means that the CNS modulates the control of the contralateral leg in an unnatural and possibly less synchonrized manner to conduct the bilateral exercise of sit-to-stand, to compensate for the reduction of natural response of the fatigued limb. It should be noted that the possibility of changes in contralateral control due to fatigue has also been acknowledged in the literature [16], [17], [45], which supports the results of the proposed network analysis. Thus, in summary it can be mentioned that this paper, for the first time illustrates the global muscle network modulation in a bilateral manner due to fatigue. The bilateral fatigue-related decreases in connectivity can be attributed to disruption of synchronicity between CNS and PNS, or the distributed control at PNS level. The bilateral effect highlights the possibility of systematic effect of fatigue on neural aspects of motor control in tasks which would require bilateral synchronicity/coordination, or it can be caused by a compensatory mechanism which modulates the neural control of the contralateral side to conduct the task.

In contrast to the consistent response from the non-parametric muscle network, conventional spectrotemporal methods (RMS, PSD and MDF) failed to detect the resistance-training induced muscle fatigue, in this study. In Fig. 6, no consistent patterns of RMS, PSD and MDF were observed for the selected muscles. Only the PSD of the right VL muscle and the MDF of the left GO muscle showed a marginally significant change, across the subject population. The increase in PSD of VL loosely corresponds to previous studies which found that myographic activation measurements (RMS) increase in response to fatigue (please note that most existing literature in this regard relates to aerobic fatigue). However, even the aforementioned changes in spectrotemporal measurements do not follow a consistent trend for all subjects. The heterogeneity of the RMS, PSD and MDF results is highlighted by the blue hairlines, which show some subjects showing an increased metric for a given muscle, and other subjects showing a decrease. High inter-subject variability within the results for the established spectrotemporal measurements in response to fatigue tallies with the existing literature [46], and emphasizes the significance of uncovering a robust biomarker of fatigue.

Regarding the broader impact of this work, it can be mentioned that accurate detection of fatigue has strong applications in physiological and rehabilitation settings. The method could be broadly used in exercise sciences, for example measuring individual muscle strength, or a muscle’s capacity to work before degradation. Additionally, the muscle group’s capability to function correctly in tandem could be monitored, since the non-parametric muscle network identifies fatigue at both the particular muscle and overall network levels. Objective quantification of fatigue would represent an improvement on current practices of self-reporting fatigue in physiotherapy [47], where accurate fatigue-tracking could help fatigue management during recovery from conditions such as cancer [48] and other functional motor disorders [49]. The proposed biomarker can help guide rehabilitation from the mentioned disorders, as well as other serious problems which lead to weaker muscles, such as a joint injury or stroke. Accurate monitoring of fatigue could help such patients and therapists to determine when the exercise limit has been reached, and prevent injury potentially caused by continuing. A further rehabilitation application of this method lies in augmented sEMG control of assistive devices. It is known that sEMG signal characteristics change in response to fatigue, and this transformation can affect sEMG-based control of prosthetics and assistive exoskeletons [50]. A precise quantification of fatigue can help such systems to accurately compensate for its effects and maintain their function.

## V. Conclusion

In this paper, a new method, namely non-parametric functional muscle network, for detecting muscle fatigue has been proposed and its efficacy has been validated. The proposed biomarker could significantly detect fatigue-related decreases at the subject group, individual subject and particular node levels. The strong performance of the non-parametric muscle network is accentuated by the heterogeneous and inconclusive response of conventional sEMG-based measurements of fatigue to the same input data. The proposed fatigue quantification technique has a broad range of applications, most significantly in physiotherapy and rehabilitation.

